# Antigen self-anchoring onto bacteriophage T5 capsid-like particles for vaccine design

**DOI:** 10.1101/2022.11.03.515007

**Authors:** Emeline Vernhes, Linda Larbi Chérif, Nicolas Ducrot, Malika Ouldali, Lena Zig, N’diaye Sidibe, Sylviane Hoos, Luis Ramirez-Chamorro, Madalena Renouard, Ombeline Rossier, Patrick England, Guy Schoehn, Pascale Boulanger, Karim Benihoud

**Affiliations:** Université Paris-Saclay, CEA, CNRS, Institute for Integrative Biology of the Cell (I2BC), 91198 Gif-sur-Yvette, France; Université Paris-Saclay, Gustave Roussy, CNRS, Metabolic and systemic aspects of oncogenesis for new therapeutic approaches (METSY), 94805 Villejuif, France; Institut Pasteur, Biophysique Moléculaire, CNRS UMR 3528, Paris, France; Univ. Grenoble Alpes, CNRS, CEA, IBS, F-38000 Grenoble, France

**Keywords:** bacteriophage T5, capsid, decoration protein, protein display, antigen, vaccine

## Abstract

The promises of vaccines based on virus-like particles stimulate demand for universal non-infectious virus-like platforms that can be efficiently grafted with large antigens. Here we harnessed the modularity and extreme affinity of the decoration protein pb10 for the capsid of bacteriophage T5. SPR experiments demonstrated that pb10 fused to mCherry or to the model antigen ovalbumin (Ova) retained picomolar affinity for DNA-free T5 capsid-like particles (T5-CLPs), while cryo-EM studies attested to the full occupancy of the 120 capsid binding sites. Mice immunisation with CLP-bound pb10-Ova chimeras elicited strong long-lasting anti-Ova humoral responses involving a large panel of isotypes, as well as CD8^+^ T cell responses, without any extrinsic adjuvant. Therefore, T5-CLP constitutes the first DNA-free bacteriophage capsid able to irreversibly display a regular array of large antigens through highly efficient chemical-free anchoring. Its ability to elicit robust immune responses paves the way for further development of this novel vaccination platform.

## Introduction

Tackling infectious endemic diseases and emerging pandemics requires new vaccines providing maximum safety, tolerability and immunogenicity. Virus-Like Particles (VLPs) offer great potential for targeted antigen (Ag) delivery and meet these requirements. They self-assemble into non-infectious particles mimicking the real virus and constitute multivalent Ag-display platforms. Their nanoscale size combined with the multimerization of the Ag on their repetitive surface geometry play a major role in their ability to trigger potent immune responses^1–3^. Currently licensed VLP-based vaccines targeting Human Papillomavirus (HPV)^4^ or Hepatitis B virus (HBV)^5^ rely on viral proteins carrying their own Ag. Yet, the need to diversify VLP-vaccines open the quest for universal virus-like scaffolds able to display heterologous Ags. Viruses infecting bacteria, or bacteriophages, have been proposed as versatile and efficient Ag-nanocarriers for mounting immune responses against different Ags^6^. Most of them enclose their genome in an icosahedral capsid that constitutes a highly stable platform for Ag-display. During the last decade, several infectious phage particles were used for vaccination assays against human pathogens in murine models^7^. Among them, bacteriophage T4 was engineered by grafting Ags to the capsid surface or by modifying the phage genome to deliver DNA encoding Ags from *Y. pestis*^8^ or more recently SARS-CoV-2^9^. While infectious phages are often easier to produce than their genome-free capsid, their use as vaccines in human clinical trials is confronted to international regulatory issues posed by the use of self-replicating viruses in medicine, as is currently the case with phage therapy^10^. In contrast, phage capsids devoid of viral genome could meet regulatory requirements applicable to VLPs. VLPs derived from RNA phages self-assemble upon expression of the gene encoding their coat protein (CP). The CP can tolerate the genetic fusion of Ag to its N-or C-terminal ends or insertion in external unstructured loops^11^. However, the self-assembly of Ag-VLP based on genetic fusion is limited to relatively small Ags (< 50 amino acids)^12^, or can be achieved if only a small proportion of CP subunits bears larger Ags^13^. Chemical crosslinking or bio-conjugation technologies, like SpyTag/SpyCatcher conjugation that creates a covalent iso-peptide bond between pre-purified VLPs and Ags, were used to overcome this limitation. However the Ag-coupling efficiency is highly variable and difficult to control, ranging from 20 to 80 % depending on the Ag^14,15^. Capsid decoration proteins that are found in some tailed bacteriophages constitute an attractive alternative to conjugation methods. These proteins spontaneously attach to their specific sites onto the mature capsid once the genome has been packaged and represent potential home bases for Ag display. The decoration protein gpD of phage lambda, which binds as trimer spikes to the three-fold axes of the capsid, was genetically or chemically modified to display heterologous Ag on self-assembled VLPs derived from Lambda capsid^16^. Although these VLPs have proven to elicit strong humoral immune responses in mice^17^, the methods used for Ag grafting do not allow full Ag load. Further development of phage capsids for vaccination requires reliable scaffolds that can efficiently anchor Ags in a precise array independently of their sizes. With this in mind, we turned our interest toward the large icosahedral capsid from bacteriophage T5 (90 nm in diameter)^18^. The capsid shell is formed of 775 subunits of the CP (the major capsid protein pb8) organized as hexamers on the faces and pentamers on 11 of the 12 vertices^19^. Its outer surface displays a monomeric decoration protein pb10 (17.3 kDa) bound at the center of each of the 120 CP hexamers^19^. pb10 is formed of an N-terminal capsid-binding domain (pN), connected by a flexible linker to a C-terminal immunoglobulin (Ig)-like domain (pC) exposed to the solvent (**Fig. 1a**). The pN domain alone anchors with the same high affinity as full-length pb10 to its binding sites (K_D_ = 10^−12^ M), while the pC domain does not interact with the capsid^20^. These properties suggest that pC could be swapped for a heterologous protein while keeping the interaction of pN with the capsid. Capsids of dsDNA bacteriophages initially assemble into compact procapsids, which undergo expansion upon genome packaging. This structural rearrangement of capsid protein subunits yields mature particles capable of withstanding the internal pressure generated by the packed dsDNA. T5 constitutes a particularly attractive system as stable empty capsids devoid of viral DNA and of decoration protein can be purified from bacteria infected with a T5 phage mutant impaired in DNA packaging. These empty capsids can be maturated in their stable expanded conformation and decorated *in vitro* with pb10^20,21^. Based on the properties of these T5 capsid-like particles (CLPs), we evaluated here their potential as multivalent vaccine platforms. Chimeric proteins composed of pb10 fused to the model Ag ovalbumin (Ova) were shown to retain high-affinity for T5 capsid, thus allowing the self-assembly of a nanoparticle displaying numerous Ag copies. Immunization of mice injected with these nanoparticles elicited robust humoral and cellular immune responses, compared to immunization with the chimeric protein alone. Our results establish the potency of T5-derived CLPs to serve as a vaccination platform without the need for extrinsic adjuvant.

**Fig. 1:**
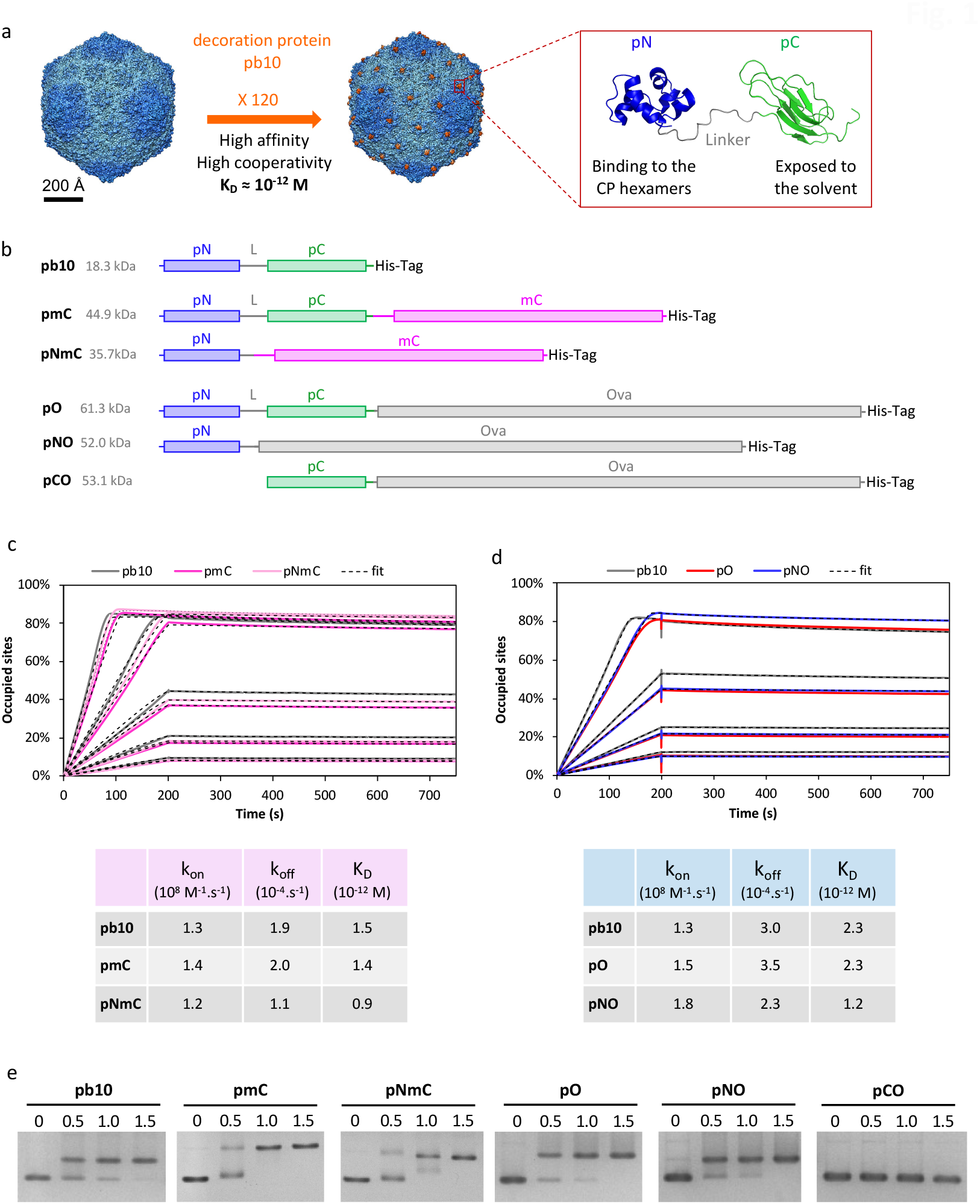
Anchoring of pb10-chimeras onto T5 CLPs with picomolar affinity. (**a**) High-affinity decoration of T5 capsid with 120 copies of the protein pb10, as seen on the surface view of T5 capsid density maps (represented from EMDB accession codes 6OMA and 6OMC). The backbone structure of pb10 shows the capsid anchoring N-terminal domain pN (blue, PDB code 5LXL) and the C-terminal domain pC (green, PDB code 5LXK). (**b**) Schematic representation of pb10-chimeras pmC, pNmC, pO, pNO and pCO used in this study, with reference to the native protein pb10. The N-terminal (pN) and C-terminal (pC) domains of pb10 are in blue and green respectively. L is the linker region between pN and pC domains. The fused heterologous proteins mCherry (mC) and Ovalbumin (O) are in pink and grey respectively. The lines represent the unfolded flexible regions while the rectangles represent the structured domains of each protein. (**c-d**) SPR real-time profiles of association with T5 CLPs (0 to 200 s) and of dissociation (200 to 750 s) using pb10 chimeras at increasing concentrations: 0.25, 0.5, 1.0, 2.0 and 4.0 nM for pb10-mC (**c**); 0.3, 0.6, 1.25, 2.5 nM for pb10-Ova (**d**). SPR response is expressed as the percentage of occupied binding-sites on the CLP, in order to cope with the mass difference of the pb10 chimeras. Below the graphs are the association and dissociation rate constants k_on_ and k_off_ calculated from the SPR profiles and the resulting dissociation equilibrium constants (K_D_). The fitting method used for the determination of these constants is described in the Methods, according to the reference^20^. (**e**) Binding assays of pb10 and chimeras to T5 CLPs analyzed by native agarose gel electrophoresis. The [pb10]/[binding site] molar ratio indicated above each lane was calculated as described in the Methods.

## RESULTS

### Protein anchoring onto bacteriophage T5 CLPs with picomolar affinity

To assess the potential of T5 CLPs as a protein display nanoparticle, we took advantage of the anchoring domain pN of the decoration protein pb10 (**Fig. 1a**)^20^. We engineered chimeric proteins formed of full-length pb10 (p) or its pN domain alone fused at their C-terminal end with a heterologous protein: either the fluorescent protein mCherry (mC, 26.8 kDa) yielding pmC and pNmC chimeras or the model Ag Ova (42.8 kDa) to form pO and pNO chimeras (**Fig. 1b**). One additional construct pCO, formed of pC domain fused to Ova, was used as a negative control for capsid decoration, as pC domain does not bind to T5 capsid^20^. The expression of all fusion genes in *E. coli* yielded soluble monomeric proteins that were purified successively by affinity, ion exchange and size exclusion chromatography as detailed in the Methods section. Protein purity was assessed by SDS-PAGE analysis (**Supplementary Fig. 1**). Binding of pb10 chimeras to T5 CLPs was first assessed by Surface Plasmon Resonance (SPR). T5 CLPs were non-covalently captured on a SPR sensor chip through anti-capsid antibodies as previously described^20^, and then association and dissociation of pmC, pNmC, pO and pNO were monitored at protein concentrations ranging from 0.25 to 2.5 or 4.0 nM. The SPR real-time profiles of association (200s) and dissociation (600s) of pb10 and its chimeras are shown in **Fig. 1c-d**. As observed for the control protein pb10, association of pmC and pNmC (**Fig. 1c**) or of pO and pNO (**Fig. 1d**) with T5 CLPs is fast, while dissociation is remarkably slow, suggesting a quasi-irreversible binding. From the determination of the association and dissociation rate constants k_on_ and k_off_ we calculated the dissociation equilibrium constants (K_D_) of 0.9 -2.3 × 10^−12^ M for the chimeric proteins, very comparable to pb10 K_D_ (1.5 -2.3 × 10^−12^ M). These values demonstrate that modification of the C-terminus of pb10 does not modify the huge affinity of the pN domain for the capsid, opening the possibility of modifying the pC domain without affecting the capsid decoration process. Binding of each pb10 chimera to T5 CLPs was also assessed by mobility shift assays in native agarose gel electrophoresis (**Fig. 1e**). CLPs appeared fully decorated with pmC, pNmC, pO and pNO proteins for a [protein]/[binding site] molar ratio in the range 1-1.5, as observed for unmodified pb10, attesting that the pN domain retains its high affinity for T5 CLP during electrophoresis, regardless of the protein linked to its C-terminal end. As expected, pCO protein did not modify T5 CLP mobility, showing that the fusion of pC with Ova Ag does not lead to unspecific binding to T5 CLP. The successful decoration of CLPs with chimeras formed of two different proteins, mCherry and ovalbumin, suggests that pb10 or pN can accommodate fusion with Ags of different sizes while maintaining their ability to irreversibly bind capsids.

We checked the integrity of the CLPs associated with pNO by cryo-electron microscopy (cryo-EM) coupled to image analysis. Representative images of T5 CLPs decorated with pNO are shown in **Fig. 2a**. Some globular extra densities are visible and decorating the surface of the capsid. They become more visible in the three-dimensional reconstruction obtained from the images. As observed for the wild-type T5 capsid structure decorated with native pb10^19^ (EMD-6OMC) (**Fig. 1a**) the pN domain is visible at low and medium contour levels (**Fig. 2b-c**, left, middle) confirming that all of the pb10 binding sites are occupied. In contrast to the wild-type T5 capsid structure, for which the pC domain of pb10 is too small and too flexible to be rendered (**Fig. 1a** and reference 19), increasing the contour level for the CLP decorated with pNO revealed some smeared densities on top of pN (**Fig. 2b-c**, right). These densities can without any doubt be attributed to Ova. The fact that Ova is fuzzy is due to the flexibility of the linker between pN and Ova, a behaviour also observed for Ags bound on the ADDOMER particle^22^. Together with the previous biochemical and kinetic analysis, cryo-EM data attests for the complete and high affinity decoration of T5 capsids with pb10 chimeras.

**Fig. 2:**
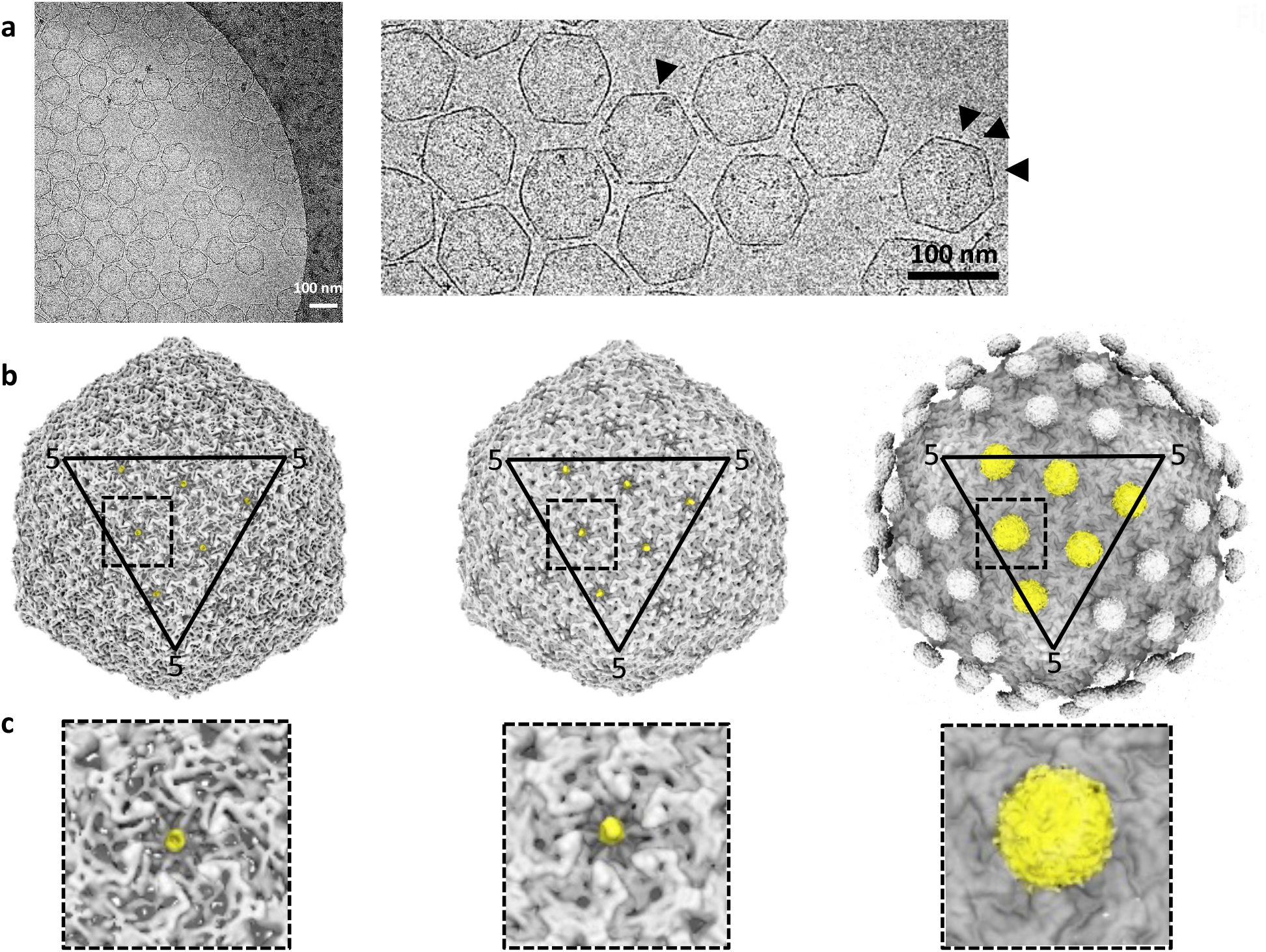
Cryo-EM image analysis and three-dimensional reconstruction of T5 CLPs decorated with pNO. (**a**) Representative example of a cryo-electron microscopy image (left) and enlarged view (right). The arrowheads indicate the presence of Ova at the surface of the CLPs. The scale bars represent 100 nm. (**b**) Isosurface representation of the 3D structure of T5 CLP decorated with pNO at low (left), medium (middle) and high (right) contour levels. One facet is highlighted by a triangle, the number 5 indicates CP pentamers at the vertices. (**c**) Enlarged views of the different squares highlighted in b and centered on a CP hexamer. Left: low contour level isosurface representation of pNO 3D structure. Even at low contour level the pN is visible (highlighted in yellow) showing that pNO occupies nearly if not 100% of the available sites in the center of hexamers. Middle: medium contour level isosurface representation of pNO 3D structure. The pN domain (yellow) is clearly visible. Right: high contour level isosurface representation of pNO 3D structure. On top of each pN domain an extra density becomes visible, which can unambiguously be attributed to Ova. The Ova density is smeared because of the presence of a flexible linker between pN and Ova: different positions of Ova are averaged out in the 3D structure.

### Humoral immune responses induced by CLP-bound pb10-Ova chimeras

Seeking to harness T5 capsid as a vaccination platform, we investigated whether anchoring of pO or pNO onto T5 CLP could modulate their immunogenicity following their administration to mice. We first checked that the endotoxin content of pb10-Ova chimeric proteins (pO and pNO) and CLP samples was in the range (20 to 200 EU/mL) acceptable for vaccine formulation^23^ **(Supplementary Table 2)**. Then, C57BL/6 mice were subcutaneously injected with pO-or pNO-decorated CLPs or with pO or pNO proteins mixed or not with complete Freund adjuvant (CFA). Sera were collected every two weeks after administration and anti-Ova antibodies (Abs) were quantified by ELISA (**Fig. 3**). The level of anti-Ova Ab (IgG) was 1000-fold higher in mice injected with either CLP-bound pb10-Ova chimeras than in mice injected with pO or pNO proteins alone. This higher titer in anti-Ova Abs was observed as soon as 14 days post-injection (p.i.) and was maintained up to 6 months (192 days p.i.). Of note, this stronger humoral immune response was observed for both pO- and pNO-CLP, suggesting that the pC domain does not impact the production of anti-Ova Abs. Remarkably, the levels of anti-Ova Abs were similar in mice injected with pO-or pNO-CLP and in mice injected with pO mixed with CFA, a compound well-known for its adjuvant properties^24^. In order to assess the quality of anti-Ova Ab responses we identified the nature of the isotypes produced at the peak of the response (day 42 p.i., **Fig. 4**). A strong production of anti-Ova Abs (*p < 0*.*001* compared to mice injected with pO or pNO alone) was observed for all isotypes analysed (IgG1, IgG2b, IgG2c and IgG3) in mice injected with CLP-bound pO or pNO. The comparison with mice injected with pO supplemented with CFA uncovered a bias towards a lower production of IgG1 (*p < 0*.*001*) and a higher production of IgG3 (*p < 0*.*05*).

**Fig. 3.**
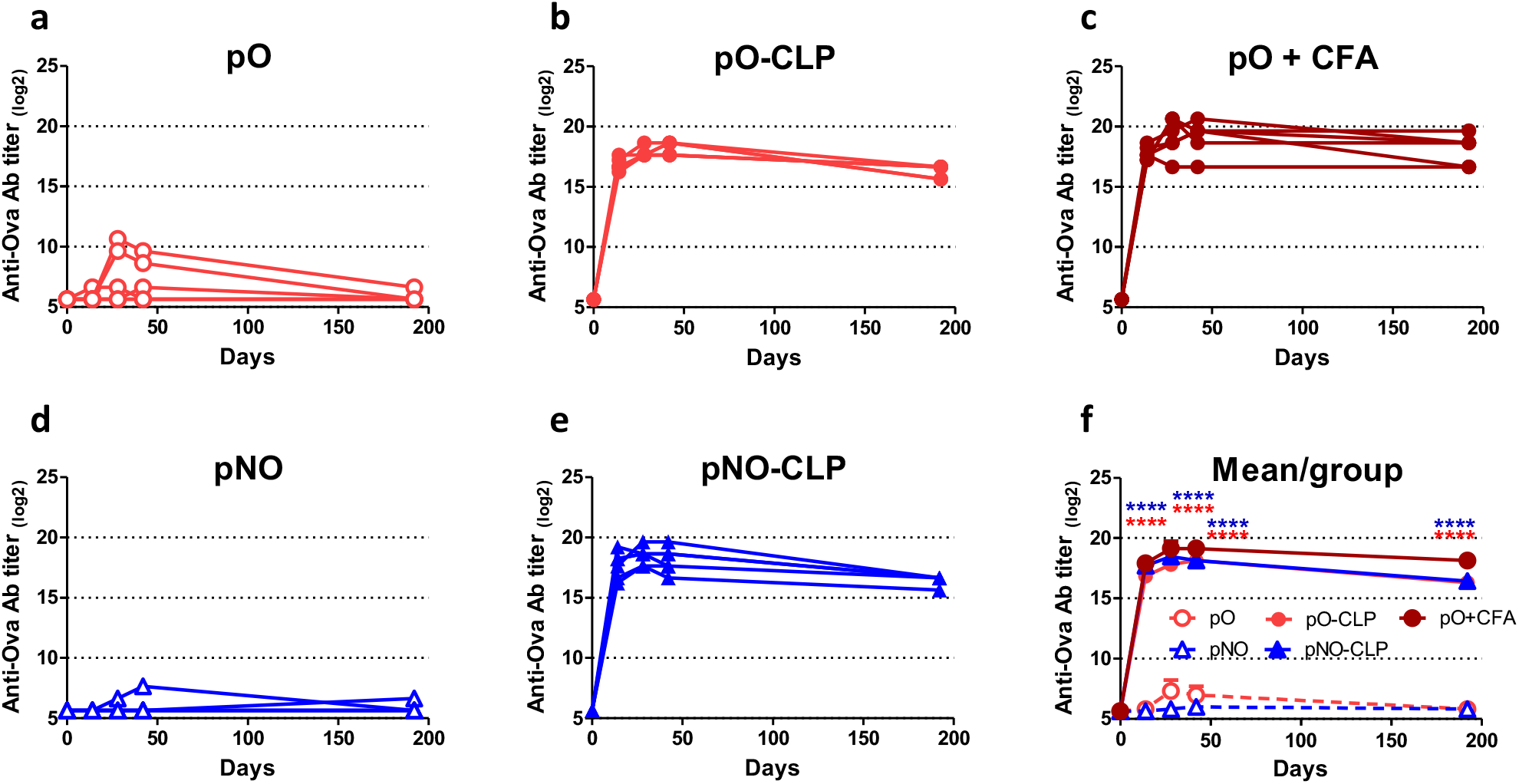
Kinetic of anti-Ova humoral immune responses elicited by pb10-Ova chimeras. C57BL/6 mice were immunized subcutaneously with pb10-Ova chimeras alone (pO or pNO), bound to T5 CLP (pO-CLP or pNO-CLP) or supplemented with CFA (pO + CFA). Titers of Ova-specific IgG were determined by ELISA at different time points after injection. (**a-e**) Each curve represents titers (log2 scale) of individual mice (n = 6) of the indicated group. (**f**) Comparison of the kinetics of the different groups (mean + SEM). Titers below 100 were plotted as log_2_(50). ****, *p < 0*.*0001* versus pO alone (red) or pNO alone (blue).

**Fig. 4.**
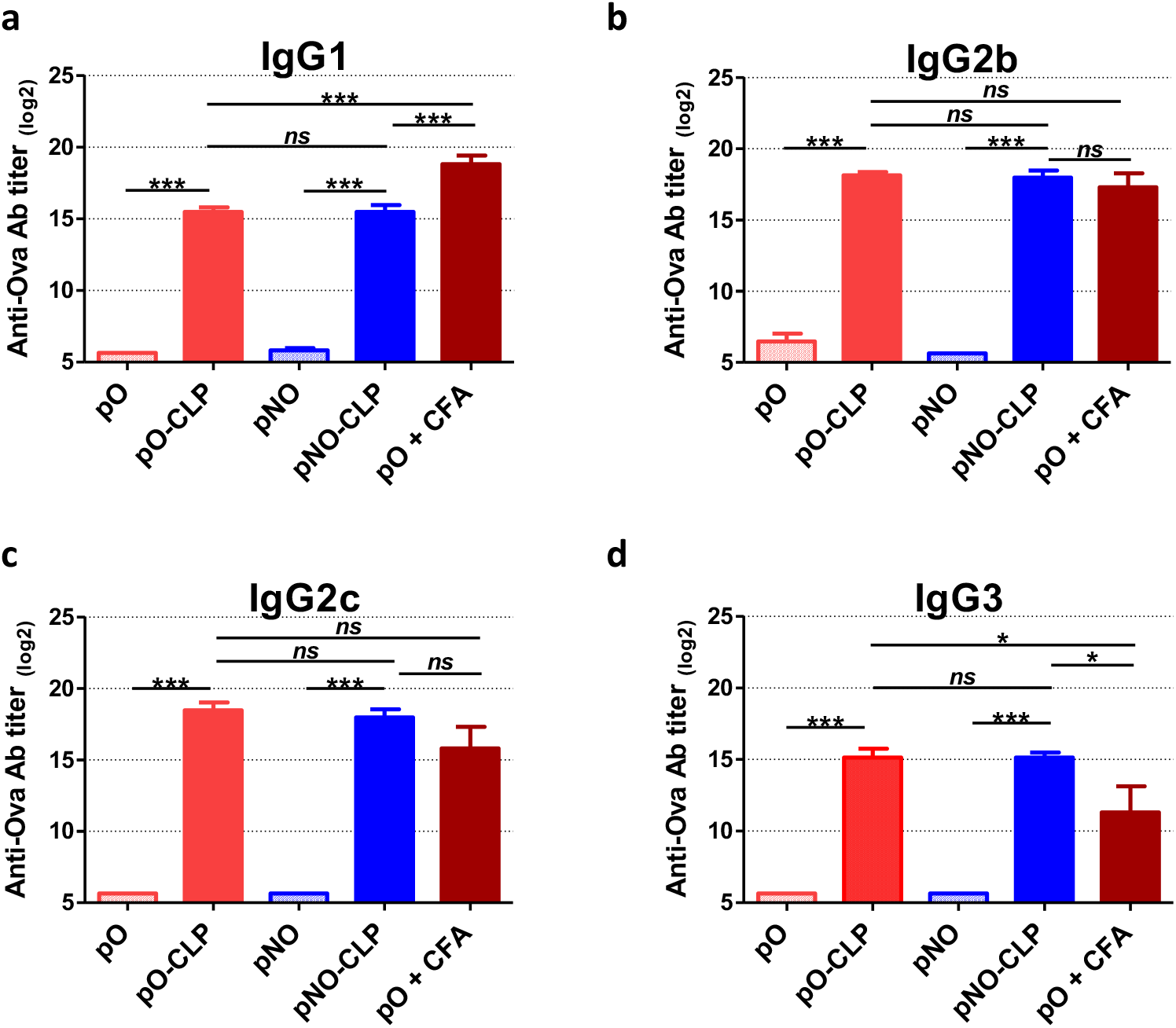
Nature of the anti-Ova humoral immune responses elicited by pb10-Ova chimeras. Mice were immunized subcutaneously with pb10-Ova chimeras alone (pO or pNO), bound to T5 CLP (pO-CLP or pNO-CLP) or supplemented with CFA (pO + CFA). Titers of anti-Ova Abs of IgG1 **(a)**, IgG2b **(b)**, IgG2c **(c)** and IgG3 **(d)** isotypes were determined by ELISA at day 42 p.i. The results correspond to the mean + SEM of each group (n = 6, log_2_ scale). Titers below 100 were plotted as log_2_(50). *ns*, non-significant; *, *p < 0*.*05*; **, *p < 0*.*01* ; ***, *p < 0*.*001*.

These data show that the binding of multiple copies of pb10-Ova chimeras to T5 CLP elicits strong and long-lasting anti-Ova humoral responses, involving a large panel of isotypes. Furthermore, since no specific adjuvant was added in the vaccine preparation, our results suggest that T5 capsid *per se* provides an adjuvant effect and influences the nature of Ova-specific Ab responses.

### Anchoring of pb10-Ova chimeras to T5 CLP is key in mounting humoral immune responses

To examine whether the attachment of the chimeric proteins to T5 CLP is mandatory to mount efficient immune responses, we used pNO and pCO chimeras, which are proficient and deficient for capsid binding, respectively (see **Fig. 1**). C57BL/6 mice were immunized subcutaneously with pNO or pCO either alone, or combined with T5 CLP, or combined with CFA. Sera were collected at different time points. As reported above, the combination of pNO with T5 CLP led to a kinetic of anti-Ova humoral responses very similar to the combination of pNO with CFA and strongly different from pNO alone (**Fig. 5a**). In sharp contrast, no significant difference in the response was observed whether pCO was injected alone or with T5 CLP as shown by the measurement of total IgGs (**Fig. 5b**) and specific isotypes (**Supplementary Fig. 2**). The weak Ab response observed with pCO mixed with capsids (pCO + CLP) did not stem from a lack of pCO immunogenicity since the combination of pCO and CFA triggered strong humoral responses (**Fig. 5b and Supplementary Fig. 2**). Altogether, these results indicate that binding of pb10-Ova chimeras to T5 CLP plays a key role in eliciting anti-Ova Ab responses. This is probably due the clustering of up to 120 copies of Ag on the same particle.

**Fig. 5.**
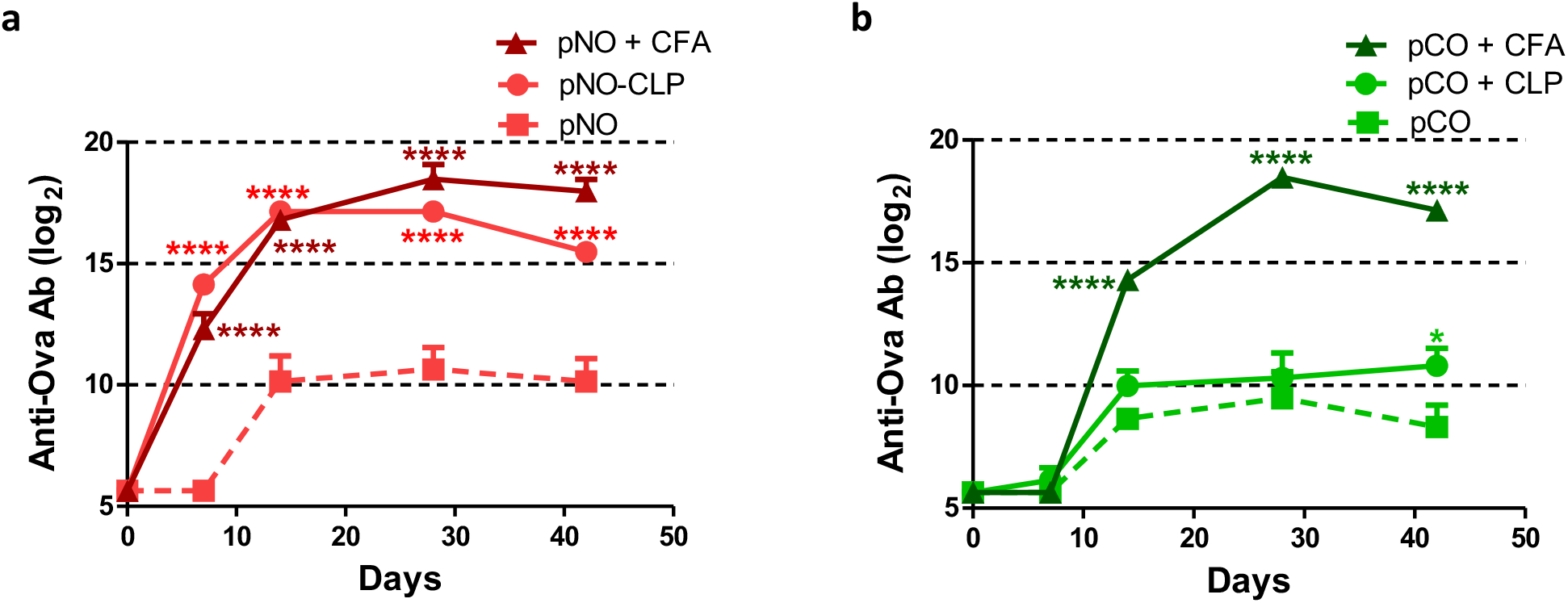
Kinetic of anti-Ova humoral immune responses elicited by Ova fused to pN or pC domains of pb10. C57BL/6 mice were immunized subcutaneously with pb10-Ova chimeras (**a**, pNO ; **b**, pCO) alone or combined with either T5 CLP (pNO-CLP, pCO + CLP) or CFA (pNO + CFA, pCO + CFA). Titers of Ova-specific IgGs were determined by ELISA at different time points after injection. The results correspond to means + SEM of data of individual mice (n = 6, log_2_ scale). Titers below 100 were plotted as log_2_(50). *, *p < 0*.*05* and ****, *p < 0*.*0001* versus pNO or pCO alone.

### Induction of strong T-cell immune responses by CLP-bound pb10-Ova chimeras

In order to assess anti-Ova cellular CD8^+^ responses elicited by administration of CLP-bound pO or pNO, splenocyte responses were quantified 10 days after boosting. ELISPOT assays showed a strong increase in IFNγ-producing CD8^+^ splenocytes in mice injected with pO-CLP or pNO-CLP compared to mice injected with pO or pNO alone (**Fig. 6a**, *p < 0*.*001* and *p < 0*.*01*, respectively). Moreover, no significant difference was observed between the CD8^+^ cellular responses elicited by pO- and pNO-CLP, suggesting that the pC domain is dispensable. Remarkably, pNO-CLP led to a significantly higher production of IFNγ-producing CD8^+^ T cells and of IFNγ than pNO combined with CFA (**Fig. 6b and 6c**, *p < 0*.*01*). Thus, in addition to their ability to induce strong humoral responses, T5 CLP displaying multiple copies of pb10-Ova chimeras constituted an efficient tool to trigger potent CD8^+^ T cell responses.

**Fig. 6.**
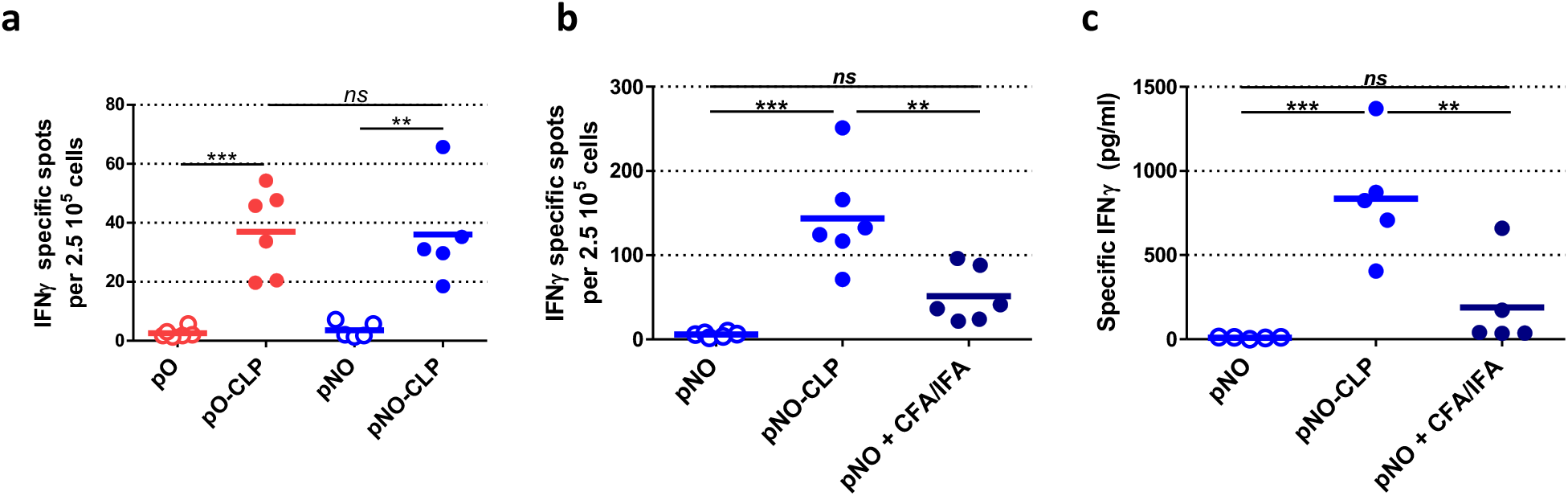
Cellular immune responses elicited by pb10-Ova chimeras. Mice were immunized (priming) with chimeric proteins either alone (pO, pNO), or bound to T5 CLP (pO-CLP, pNO-CLP) or supplemented with CFA. Mice were subsequently boosted under the same conditions (except that CFA was replaced by incomplete Freund adjuvant, ICF). **(a-b)** IFNγ-producing splenocytes were quantified by ELISPOT and (**c**) IFNγ production by splenocytes was measured by ELISA after *in vitro* restimulation with Ova_257-264_ peptide 10 days after boosting. The interval between priming and boosting was of two (**a**) or six months (**b**). The bars correspond to the mean of each group (n = 5-6) and the circles to results of individual mice. *ns*, non-significant; **, *p < 0*.*01* ; ***, *p < 0*.*001*.

## Discussion

The capsid of bacteriophage T5 offers major advantages for Ag display and vaccination. The procapsid form, devoid of DNA, is easy to produce and to maturate into a capsid-like particle (CLP) with the same structure, stability and affinity for the decoration protein pb10 compared to the native virion^20,21^. In this study, we demonstrated that the fusion of mCherry (25.8 kDa) or ovalbumin (42.8 kDa) to full-length pb10 (17.3 kDa) or to its capsid binding domain pN (8.1 kDa) yielded soluble chimeric proteins retaining picomolar affinity to T5 capsid. Such quasi-infinite affinity ensures the full occupancy of capsid binding sites (120 copies) as evidenced by cryo-electron microscopy data. The easy and regular anchoring of pb10-chimeras onto T5 CLP by mere molecular recognition between pN and CLP is better controlled and more efficient than chemical or bio-conjugation technologies. These remarkable T5 CLP properties are instrumental in achieving the production of a new nanoparticle displaying large proteins, including Ags of interest.

We probed the vaccine properties of T5 CLPs displaying pb10-Ova chimeric proteins pO or pNO. Their administration to mice elicited long-lasting anti-Ova Ab responses as well as CD8^+^ T cell responses. A single injection of non-adjuvanted CLPs decorated with pO or pNO was sufficient to elicit a strong production of anti-Ova Abs for more than 6 months. Remarkably, these responses were similar to the ones obtained after co-administration of pO or pNO with CFA. The characterization of Ab responses revealed the production of a large panel of isotypes (IgG1, IgG2b, IgG2C and IgG3). This suggest the capacity of Ag-decorated T5 CLP to mobilize different subpopulations of helper or/and follicular helper T cells (Th/Tfh) able to promote B-cell differentiation. The production of different isotypes is of major interest to activate different Ab effector functions such as neutralization, phagocytosis, complement activation and cell cytotoxicity^25^. Such large isotypic responses were not reported previously upon vaccination using bacteriophage particles, neither with the infectious bacteriophage T4 displaying Ags from *Yersinia pestis*^26^ nor with PP7-derived VLP displaying HPV epitopes^27^. Beside the humoral responses, we demonstrated that CLPs displaying pb10-Ova proteins elicited a strong induction of CD8^+^ T cell responses. In previous studies, Pouyenfard *et al*. reported that T7 bacteriophages displaying a CD8^+^ T cell epitope derived from a tumor Ag were able to trigger potent anti-epitope T cell responses^28^. Also, T4 bacteriophage heads (capsids filled with viral DNA) displaying an Ag from *Y. pestis* were shown to induce CD8^+^ T cell responses^29^. To the best of our knowledge, this study is the first to report induction of both humoral and cellular responses by a fully-decorated capsid shell devoid of viral DNA.

Vaccination studies using infectious bacteriophages (λ^30,31^, PP7^27^, T4^26^ or T7^28^) or phage heads (Qβ^32^ or T4^29^ containing RNA or DNA respectively) reported the induction of immune responses in the absence of adjuvant. Similarly, complexes between T5 CLP and pb10-Ova chimeras triggered potent humoral and cellular responses in absence of any extrinsic adjuvant. As T5 CLP preparations contain low amounts of endotoxin, our results suggest that T5 capsid shell alone, in the absence of phage genome, provides the adjuvant effect required for efficient vaccination. We speculate that the capsid itself or some structural motifs might be detected by Ag-presenting cells as previously documented for different viruses^33–35^. The strong immunogenicity of T5 CLPs may stem also from the highly ordered and repetitive distribution of Ags (120 copies), which is known to promote the engagement of several B-cell receptors at the surface of Ag-specific B-cells^1,36^. This immunogenicity might also be favoured by the delivery of Ag-CLP complexes to B-cell follicles within secondary lymphoid organs, as reported for VLPs derived from other bacteriophages^37.^

This study paves the way for the development of T5 CLP-based nanoparticles as a new platform for Ag delivery. Their attractiveness relies on several key features: (i) the tremendous thermal stability of T5 CLPs, which resists temperatures up to 95°C^21^; (ii) the easiness and low-cost of their large scale production in *E. coli* cells; (iii) the intrinsic adjuvant properties of T5 CLPs, (iv) the capacity of the decoration protein pb10 or its pN domain alone to tolerate the fusion with large size Ags and finally (v) the well-controlled and highly efficient anchoring of a high copy number of the displayed Ag. Further investigations will establish the potential of this platform to protect against different pathogens.

## Supporting information

Vernhes et al. Supplemental Information

## Methods

### Expression vectors for the production of pb10 chimeric proteins

The coding sequence of the full-length decoration protein pb10 (GenBank accession number: AAU05286) was previously cloned into the pET28b vector in frame with a C-terminal His-Tag^20^. The gene encoding variants of pb10 fused in their C-terminal end with mCherry (Red fluorescent protein, GenBank accession number AAV52164) or Ova (Ovalbumin, GenBank accession number J00895) were cloned in pET28b: (i) full-length pb10 for pmC and pO, (ii) the N-terminal capsid binding domain of pb10 (pN, first 73 amino-acids) for pNmC or pNO and (iii) the C-terminal domain of pb10 (pC, amino acids 74 to 164) for pCO. All constructs included a His-Tag in frame with the mCherry or Ova C-terminal end. The pET28-pmC, -pNmC, -pO and -pNO vectors were constructed using a ligation-free technology (FastCloning method according to Li et al^38^ or with the Quick-Fusion Cloning Kit from Biotool), while pET28-pCO was constructed by using the golden-gate seamless assembly method based on the type IIS restriction endonuclease according to Engler *et al*.^39^. The cloning strategies and oligonucleotides used for generating these expression vectors are detailed in supplementary information and **Supplementary Table 1** respectively.

### Production and purification of pb10 chimeric proteins

*E. coli* BL21 (DE3) cells harbouring each of the different pb10 expression vectors were grown in LB medium supplemented with 50 μg/mL kanamycin at 37°C for pmC, pNmC, and pCO or at 28°C for pO and pNO (increased solubility). At mid-exponential growth phase (OD_600nm_ = 0.6 -0.8), protein expression was induced by addition of 0.4 mM isopropyl-β-D-thiogalactopyranoside (IPTG) and the growth continued for 2-3h. Bacterial cells harvested by centrifugation were suspended in the loading buffer (50mM Tris-HCl pH 7.4 containing 1M NaCl), broken by two passages in a French press (10,000 psi) at 4 °C and centrifuged at 4 °C (100,000 g, 30 min). The supernatant was incubated at 4 °C for at least 8 hours with 1% n-Octyl-β-d-Glucopyranoside (OG), a detergent used to solubilize contaminant endotoxins (Lipopolysaccharide molecules) originating from *E. coli* outer membranes. It was then loaded onto a 5 mL HisTrap™ FF column (Cytiva) pre-equilibrated in the loading buffer supplemented with 0.2% Lauryldimethylamine-N-oxide (LDAO) and connected to an ÄKTA purifying system. The pb10-mC or -Ova chimeras were eluted with a 0-1 M imidazole gradient in the presence of 0.1% LDAO and the eluted fractions were collected for further purification in the absence of detergent, by cation or anion exchange chromatography, depending on the calculated isoelectric point (pI) of the pb10 fusion proteins. Full-length pb10 (pI = 7.9) was purified on a 5 mL HiTrap SP column (Cytiva) as previously described^20^, while pmC (pI = 6.4), pNmC (pI = 7,0), pO (pI = 6.0), pNO (pI = 6.2) and pCO (pI = 5.4) were purified on a 5 mL HiTrap Q HP column (Cytiva) pre-equilibrated in 50 mM Tris-HCl buffer pH 8.0. The proteins were eluted with a 0-1 M NaCl gradient, concentrated on a centrifugal filter (Amicon® Ultra-4, 10 kD, Millipore) and finally purified by size exclusion chromatography on a Superdex 75 10/300 column (Cytiva) pre-equilibrated in Phosphate-Buffered Saline (PBS). Protein concentrations were determined by measuring the absorbance at 280 nm and using the theoretical extinction coefficients of 22,920 M^−1^cm^−1^ for pb10, 57,300 M^−1^cm^−1^ for pmC, 45,840 M^−1^cm^−1^ for pNmC, 54,320 M^−1^cm^−1^ for pO and 42,860 M^−1^cm^−1^ for pNO and pCO determined with the online Expasy ProtParam tool^40^.

### Production and purification of T5 CLPs

In order to produce phage T5 empty capsids (CLPs) lacking pb10, we constructed the double mutant T5ΔdecstAmN5, by cross-infection of the suppressive *E. coli* strain CR63 with the mutant T5stAmN5 bearing an amber mutation in the terminase gene (production of a non-functional truncated terminase in a non-suppressive strain, thus preventing DNA packaging^18^) and the mutant T5Δdec deleted in the gene encoding the decoration protein pb10 (see ref. ^20^ for the detailed procedure of mutant screening). T5 CLPs were produced by infection of non-suppressive *E. coli* strain F with the double mutant T5ΔdecstAmN5 and purified as previously described^21^ with some modifications in the protocol to remove contaminant endotoxins. Briefly, after precipitation of the bacterial lysate with polyethylene glycol followed by centrifugation in glycerol gradients, the fractions containing T5 empty capsids were incubated for at least 8 hours with 1% of OG detergent. Then, a first step of anion-exchange chromatography was performed on HiTrapQ HP column (Cytiva) equilibrated in 50 mM Tris buffer, pH 8.0 containing 200 mM NaCl and 0.2% LDAO. After their elution with a 0-1 M NaCl gradient, the empty capsids were dialyzed against 50 mM Tris buffer, pH 8.0 containing 200 mM NaCl without LDAO and re-injected onto the HiTrapQ HP column for a second step of purification by using the same binding and elution solutions without detergent. Transition of T5 empty capsid from their compact state to their stabilized expanded conformation (CLP) was obtained by dialysis against 50 mM Hepes buffer, pH 7.0, for 24-48 h^21^. CLP samples were finally dialyzed against PBS, concentrated on a centrifugal Amicon® Ultra-4 100K filter unit (Millipore) and stored at 4 °C. CLP concentration was calculated from the measurement of protein concentration as described previously^20^.

### Determination of endotoxin content

Endotoxin levels in fusion protein and CLP samples used for *in vivo* experiments were quantified using the Pierce LAL Chromogenic Endotoxin Quantitation Kit according to manufacturer’s instructions (Thermo Fisher Scientific). The results are expressed as endotoxin unit (EU) per ml.

### Surface Plasmon Resonance binding Assay

SPR experiments were conducted at 25 °C on a T200 instrument (GE Healthcare) using a CM5 sensor chip functionalized by covalent amine coupling of rabbit antibodies raised against empty T5 capsids as described in ref.^20^. The CLPs (0.2 mg/mL) were captured by injection at 5 μL/min in running buffer (PBS with 0.05 % Tween 20 and 1 mg/mL BSA) yielding a capsid density of about 900 RU (**Supplementary Fig. 3**). Native pb10 or its chimeric forms (diluted in running buffer to 0.25 -4 nM) were injected for 200 s at 50 μL/min. Running buffer was then flowed for 550 to 700 s at 50 μL/min to monitor protein dissociation. The functionalized surface was then regenerated by 0.85 % phosphoric acid for the next cycle of CLP capture followed by protein association and dissociation. SPR sensorgrams were corrected for non-specific pb10 binding to the anti-capsid surface and for buffer effect by subtracting both pb10 responses on the functionalized surface without CLPs and buffer response on captured CLPs. Kinetic evaluation was performed by fitting the experimental curves to a simple Langmuir model using the Biacore T200 Kinetics Summary Software (version 2.0, GE Healthcare). The percentage of occupied binding sites was calculated by dividing the protein association response expressed in RU by the expected response for total capsid decoration (R_100%_) determined by the following formula: R_100%_ = R_CLP_ * 120 * MW_protein_ / 26,018,181 (with R_CLP_ being the captured CLP response, 120 the number of pb10 binding sites per CLP, MW_protein_ the molecular weight of the pb10-chimeras and 26,018,181 Da the molecular weight of an empty capsid^41^).

### Binding assays assessed by native agarose gel electrophoresis

Purified T5 CLPs were mixed with various amounts of the different pb10 constructs at a final capsid concentration of 10 nM and incubated at 4°C for 30 min. The samples were loaded on a 1.5% agarose gel in TAMg buffer (40 mM Tris-HCl, 20 mM acetic acid, 1 mM MgSO_4_, pH 8.1) and migrated at 25 V overnight in a cold room. The capsid bands were stained with Coomassie blue. The molar concentration of binding sites was calculated by multiplying the capsid concentration by 120 sites per capsid.

### Cryo-EM, image analyses and 3D reconstruction

A 3.5 μL sample of concentrated pNO-decorated CLPs was applied to negatively glow discharged (25 mA, 40 s) R3.5/1 quantifoil copper grids (Quantifoil Micro Tools). The excess of solution was blotted using a Vitrobot Mark IV (FEI) (20 °C, 100 % humidity, 2 s blot-ting time and blot force 1) and subsequently flash-frozen in liquid ethane. Automated data collection was performed on a 200 kV Glacios cryo-TEM microscope (Thermo Fischer Scientific) equipped with a K2 direct electron detector (Gatan) using SerialEM^42^. Coma and astigmatism corrections were also performed using SerialEM. Movies of 40 frames were recorded in counting mode at a 36,000× magnification giving a pixel size of 1.145 Å with defocus ranging from −1.0 to −2.5 μm using a multi-shot scheme (3×3 grids of holes without moving the stage). Total exposure dose per movie was 40 e−/Å^2^ and total number of images was 2500.

Movie drift correction and CTF determination were performed with Relion^43^. A total number of 12,000 T5 CLPs were automatically selected into 1024 × 1024 pixels boxes from the best 500 images. These boxes were rescaled to 420 × 420 pixels boxes (pixel size of 2.9 Å) and submitted to 2D classification. After extensive selection and generation of an initial model imposing I4 symmetry, 3D refinement generated a final reconstruction including 10,449 particles with a resolution of 5.8 Å (Fourier Shell = 0.143, not shown).

### Mouse immunization

Six-week-old C57BL/6 female mice were purchased from Janvier (Le Genest Saint Isle, France). All mice were conditioned for at least 1 week in our animal facilities before beginning the experiments. All animal experiments were approved (authorization number 19055-2919021108472030 v3) by Ethics Committee No. 26 (officially recognized by the French Ministry for Research) in accordance with the European Directive 2010/63 UE and its transposition into French Law.

Recombinant fusion proteins (3.8 μM) incubated or not with T5 CLP (32 +/-3 nM, counting as 3.8 μM of binding sites) were injected subcutaneously in a volume of 50μl of phosphate-buffered saline (PBS). For each immunization, the total amount of injected proteins was 41 μg of CLPs, decorated with 11.6 μg or 9.9 μg of pO or pNO respectively, or mixed with 10.1 μg of pCO. In some experiments, a control group received injection of recombinant fusion proteins (3.8 μM) mixed with complete Freund adjuvant (CFA, Sigma) in 50 μl of PBS. Similar conditions were used when a second injection was performed, except for the control group for which recombinant fusion proteins were mixed with incomplete Freund adjuvant (IFA, Sigma). Blood samples were collected from the submandibular vein at different time points for more than 6 months and sera were prepared and analysed for the presence of specific antibodies by ELISA as described below. Spleens were collected 10 days after the second injection to monitor cellular immune responses.

### Measurement of humoral immune responses

After coating 96-well plates (Nunc) with 1μg ovalbumin (Sigma), serial dilutions of the sera in 5% milk PBS-Tween 0.05% were added. Bound antibodies were detected with peroxidase-conjugated anti-mouse IgM, IgG, IgG1, IgG2b, IgG2c or IgG3 isotype goat Abs (Southern Biotechnology Associates). The peroxidase activity was revealed by incubation with the substrate O-phenylenediamine dihydrochloride (Sigma–Aldrich) for 30 min. The reaction was stopped by addition of 3N HCl and spectrophotometric readings were performed at 490 nm. Titers were defined as the reciprocal of the highest dilution giving an OD_490_ 2-fold above background values.

### Measurement of cellular immune responses

Spleens were crushed in RPMI medium with 5 % foetal calf serum and 5 × 10^−5^ M β-mercaptoethanol, and filtered through a 100 μm cell strainer. After removal of blood cells by ACK Lysing Buffer (Invitrogen), the cells were resuspended and the concentration was adjusted at 2.5 × 10^6^ cells/mL. Then splenocytes were restimulated in different conditions (medium alone, Ova_257-264_ peptide (5μg/mL)) in a volume of 200μL for one day (ELISPOT) or three days (ELISA). For ELISPOT, plates were revealed with supplied reagents (murine IFNγ ELISPOT kit, Diaclone) and spots were counted with the ImmunoSpot® S6 FluoroSpot Line Plate Reader (C.T.L.). For ELISA, the supernatants were collected and assayed for the presence of IFN-γ using a murine IFN-γ kit (eBioscience). In both ELISPOT and ELISA, restimulation with ionomycine (1μM) and PMA (0.1μM) was used to control the viability of lymphocytes.

### Statistical Analysis

Data from ELISA experiments (titers) were log_2_-transformed before analysis. Two-way repeated measures ANOVA was used for comparison of responses measured for different groups at different time points, then Bonferroni post hoc test was used to compare between groups at each time point. Data obtained from splenocyte restimulation assays were analyzed by a one-way ANOVA followed by Tukey’s post-hoc test to compare sets of data. All graphs and statistical tests were performed using GraphPad Prism software. Differences were considered significant when *p < 0*.*05*.

## Author contributions

P.B. and K.B. designed and led the study. E.V. constructed the expression vectors for most pb10-mC and -Ova chimeras, produced and purified the proteins. E.V. and M.R. purified T5 CLPs, performed the decoration assays, and prepared the samples for mice immunization. E.V. and S.H. performed the SPR experiments and analyzed the data with the help of P.E. L.R.-C. and O.R. designed the expression vector for pCO production. N.D. prepared the CLPs decorated with pNO for EM imaging, M.O. performed preliminary negative-stain and cryo-EM imaging to define the best conditions for pNO-CLP reconstructions. G.S. performed capsid imaging on the GLACIOS microscope and 3D reconstruction of pNO-CLP. L. L.C. measured endotoxin content of protein and CLP samples, performed mice immunizations, sampled blood and spleen to monitor immune responses using ELISA and ELISPOT. L.Z. and N.S. performed ELISA experiments. P.B. and K.B. wrote the manuscript with contributions of E.V., O.R. and G.S. All authors reviewed the manuscript.

## Acknowledgements

This work was supported by the Centre National de la Recherche Scientifique and the University Paris-Saclay. L. L.C., E.V. and N.D. received a fellowship from the French Ministère de l’Enseignement Supérieur et de la Recherche. L. L.C. received an additional fellowship from the Ligue Nationale contre le cancer. L.R.-C received a fellowship “Becas Don Carlos Antonio Lopez” from the Paraguayan Ministry of Education. This work benefited from the Cryo-EM platform of I2BC, supported by the French Infrastructure for Integrated Structural Biology (FRISBI) [ANR-10-INSB-05-05] and member of IBISA. This work used the platforms of the Grenoble Instruct-ERIC center (ISBG; UAR3518 CNRS-CEA-UGA-EMBL) within the Grenoble Partnership for Structural Biology (PSB), supported by FRISBI (ANR-10-INSB-05-02 & project ID 160 to P. B.) and GRAL, financed within the University Grenoble Alpes graduate school (Écoles Universitaires de Recherche) CBH-EUR-GS (ANR-17-EURE-0003). The electron microscope facility is supported by the Auvergne-Rhône-Alpes Region, the Fondation pour la Recherche Médicale (FRM), the fonds FEDER and the GIS-Infrastructures en Biologie Santé et Agronomie (IBiSA). IBS acknowledges integration into the Interdisciplinary Research Institute of Grenoble (IRIG, CEA). The authors are very grateful to the staff of the animal facility of Institut Gustave Roussy for technical help. We thank Didier Poncet and Pierre Bobé for their critical reading of the manuscript.

## Data availability

The data supporting the conclusions of the study are available from the corresponding authors. The EM map generated in this study has been deposited in the Electron Microscopy Data Bank under the number EMD-14863.

## Declaration of interests

P.B., E.V., N.D., L. L.C., and K.B are named as inventors on a patent application filed by the CNRS and University Paris-Saclay based on the presented study (N° PCT/FR2020/051628, WO2021053309).

## Notes

### Competing Interest Statement

Funding sources:
Ministere de l'Enseignement Superieur, de la Recherche, de la Science et de la Technologie (MESRST) - ED 569 To E. Vernhes
Ministere de l'Enseignement Superieur, de la Recherche, de la Science et de la Technologie (MESRST) - ED 582 to Linda Larbi Cherif
Ministere de l'Enseignement Superieur, de la Recherche, de la Science et de la Technologie (MESRST) - ED 569 to N. Ducrot
"Becas Don Carlos Antonio Lopez", Paraguayan Ministry of Education to L. Ramirez-Chamorro

