## Supplementary material for "Antigen self-anchoring onto bacteriophage T5 capsid-like particles for vaccine design": Vernhes et al. Supplemental Information

### SUPPLEMENTARY INFORMATION

#### Construction of the expression vectors for pb10-mCherry and pb10-Ovalbumin chimeras

##### Plasmids pET28-pmC, pET28-pNmC, pET28-pO and pET28-pNO

The expression vectors encoding the fusion proteins were constructed using a ligation independent PCR cloning technology (FastCloning method according to Li *et al.*[38] or Quick-Fusion Cloning Kit from Biotool).

For building pb10-mCherry chimeras, the coding sequence of pb10 or its capsid binding domain (pN) were cloned into a pET28b vector encoding mCherry-6His (pET28-mC, a kind gift from Ahmed Bouhss, INSERM U1204, Université Paris-Saclay). The pET28-mC vector was amplified by PCR using a forward primer containing the mCherry gene's first 6 codons and a reverse primer complementary to the NcoI-RBS sequence. The sequences of pb10 or pN inserts were PCR amplified from pET28-pb10 vector[20], using primers including extensions complementary to the ends of pET28-mC PCR product. The two purified amplicons were mixed together according to the FastCloning method[38] before transformation of XL1-Blue competent cells (Agilent Technologies).

For building pO and pNO chimeras, the pET28-pb10[20] vector was amplified by PCR using a forward primer containing the 6His-Tag coding sequence and a reverse primer overlapping the last codons of pb10 sequence (for pO) or the sequence coding the linker that separates the two domains of pb10 (for pNO). The Ova insert coding sequence was PCR amplified from the pcDNA3-OVA plasmid (Addgene). We used a forward primer containing an extension overlapping the last codons of pb10 gene (for pO) or of the linker (for pNO) and a reverse primer extended with the 6His-Tag coding sequence of pET28-pb10 vector. The purified Ova PCR products were mixed with their respective pET28-pb10 PCR products and incubated with the Fusion Enzyme according to the Quick-Fusion Cloning Kit (Biotool) protocol before transformation of XL1-Blue competent cells (Agilent Technologies).

The oligonucleotides used for the above constructions are detailed in **Supplementary Table 1**.

**Plasmid pET28-pCO**

The fusion of Ova coding sequence with pb10 C-terminal domain (pC) was obtained by means of the Golden Gate cloning method, using type II restriction endonuclease BsaI that cuts DNA outside of its recognition site[39]. We amplified by PCR the two following fragments: i) the insert sequence encoding the pC-Ova region (pCO) from pET28-pO vector (see above); ii) the recipient vector pET28b including a 6His sequence to be fused with pCO C-terminal end. For these PCR reactions, we used forward and reverse primers complementary to the region to be amplified at their 3' end and flanked with a BsaI site at their 5' end, such that digestion of the fragments removes the enzyme recognition sites and generates ends with complementary four nucleotides overhangs. The cloning step was performed using a one-step restriction-ligation as follows. The amplicons were purified (GeneJet PCR Purification Kit, Thermo Scientific) and 200 ng of each product was added to a mix containing 1U of T4 ligase (Thermo #EL0011), 10 U of BsaI (NEB #R0535), 1 µL DpnI (Thermo #FD1703) in 1X T4 ligase buffer. The mix was incubated for 60 cycles of 37 °C and 22 °C, 5 minutes each, followed by the final steps at 50 °C for 1 hour and 98 °C for 20 min. The assembly reaction was then used for transformation of XL1-Blue competent cells (Agilent Technologies). The oligonucleotides used for pET28-pCO construction are detailed in **Supplementary Table 1**.

The sequence of each construction were checked by Sanger DNA sequencing.

### Supplementary Table 1

#### Oligonucleotides and PCR products used for the construction of expression vectors encoding pb10-mCherry and pb10-Ova chimeras.

| *PCR product* | **Oligonucleotides** | | **Vector** |
|---|---|---|---|
| | **Forward Primer* | **Reverse Primer* | |
| pET28-mC vector | ATGGTGAGCAAGGGCGAG | CATGGTATATCTCCTTCTTAAAGT | |
| pb10 insert | GAAGGAGATATACCATGGGGATTG | GCCCTTGCTCACCATGCCTCCAGGAACAGTAGG | **pET28-pmC** |
| pN insert | GAAGGAGATATACCATGGGGATTG | GCCCTTGCTCACCATTGGTGGAGCCGGTGGGG | **pET28-pNmC** |
| pET28-pb10 vector | CACCACCACCACCACCACTGAG | GCCTCCAGGAACAGTAGGATTAAC | **pET28-pO** |
| Ova insert | CCTACTGTTCCTGGAGGCATGGGCTCCATCGGC | GGTGGTGGTGGTGGTGAGGGGAAACACATCTGC | |
| pET28-pN vector | CACCACCACCACCACCACTGAG | TGGTGGAGCCGGTGGGG | **pET28-pNO** |
| Ova insert | CCACCGGCTCCACCAATGGGCTCCATCGGC | GGTGGTGGTGGTGGTGAGGGGAAACACATCTGC | |
| pET28-6His vector | AAAA**GGTCTC**A⇣GAATTATCTCCTTCTTAAAGTTAAACAAAATTATTTC | AAAA**GGTCTC**A⇣CACCACCACCACCACCACTGAG | **pET28-pCO** |
| pCO insert | AAAA**GGTCTC**A⇣ATTCATGTTAACTCTTTCTAAAGATTTAACTGCTAGC | AAAA**GGTCT**CA⇣GGTGAGGGGAAACACATCTGCCAAAG | |

In Bold: BsaI recognition sequence. Underlined bases indicate the 4 nucleotides overhangs generated by BsaI cleavage at N1 position (⇣) after the recognition sequence.

*Primers are written in the 5' to 3' orientation.

### Supplementary Figure 1

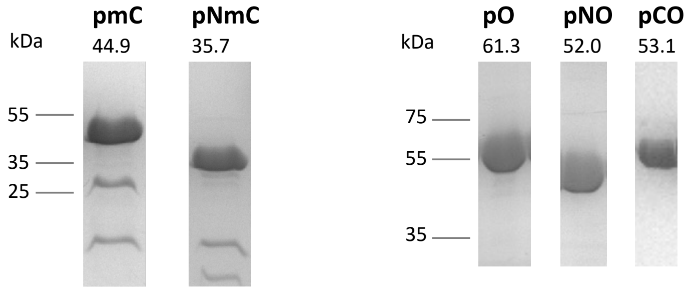


**Production of the p10 chimeras.** SDS-PAGE analysis of purified pb10-mC and pb10-Ova chimeric proteins eluted from a Superdex 75 10/300 gel filtration column. It should be noted that the bands observed below pmC and pNmC on the gel (left) are not contaminant proteins but results from the slow degradation pmC and pNmC during their storage.

### Supplementary Table 2

**Endotoxin and protein content of CLP and pb10-Ova chimera doses administered to mice**

| | **CLP** | | | **pb10-Ova chimera** | | | | **Injected dose** | | |
|---|---|---|---|---|---|---|---|---|---|---|
| **Group** | µg/mL | EU/mL | EU/100 µg | | µg/mL | EU/mL | EU/100µg | µg/mL | EU/mL | EU/100µg |
| ***pO-CLP** | 832 | **9.4** | 1.1 | *pO* | 235.2 | **4.6** | 2.0 | 1067.2 | **14.0** | 3.1 |
| ***pNO-CLP** | 832 | **9.4** | 1.1 | *pNO* | 200 | **8.4** | 4.2 | 1032.0 | **17.8** | 5.3 |
| ****pNO-CLP** | 832 | **23.6** | 2.8 | *pNO* | 200 | **18.5** | 9.3 | 1032.0 | **42.1** | 12.1 |
| ****pCO + CLP** | 832 | **23.6** | 2.8 | *pCO* | 204 | **0.6** | 0.3 | 1036.0 | **24.2** | 3.1 |

EU, Endotoxin Unit
* Samples used to obtain immunization data presented in Figures 3, 4 and 6a
** Samples used to obtain immunization data presented in Figures 5 and 6b as well as Supplementary Figure 2

**Supplementary Figure 2**

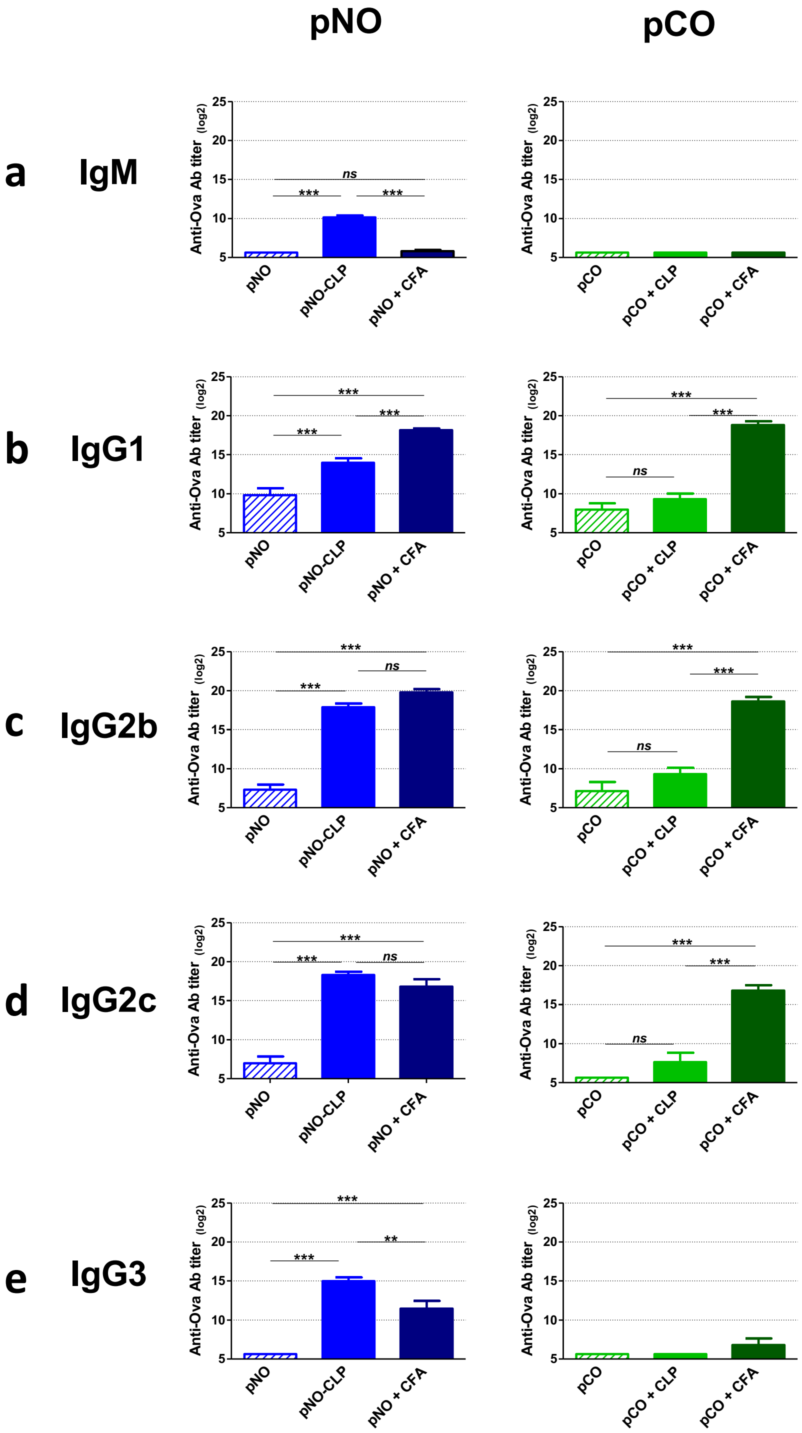


**Isotype responses of anti-Ova antibodies elicited by pNO and pCO chimeras.** Mice were immunized subcutaneously with pb10 chimeric proteins (**Left**, pNO; **Right**, pCO) alone or in combination with either T5 CLPs or CFA. Titers of anti-Ova IgM (**a**), IgG1 (**b**), IgG2b (**c**), IgG2c (**d**) and IgG3 (**e**) Ab isotypes were determined by ELISA at day 42 p.i. The results correspond to the mean + SEM of each group (n = 6, $log_2$ scale). Titers below 100 were plotted as $log_2(50)$. *ns, non-significant; **, p < 0.01 ; ***, p < 0.001.*

**Supplementary Figure 3**

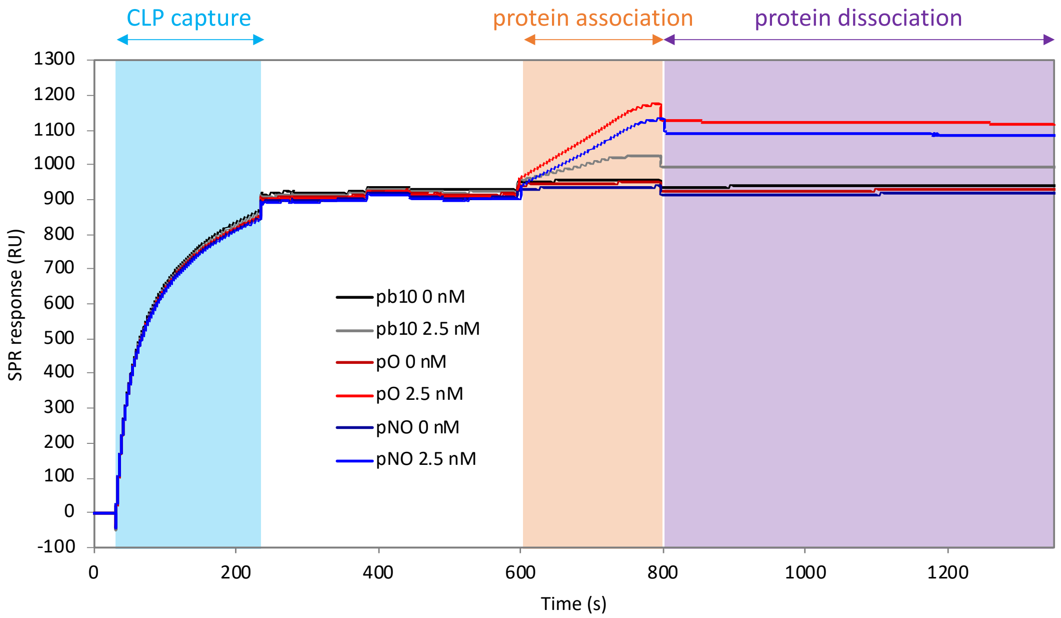


**Typical SRP profiles of CLP capture (200 s) followed by association (200 s) and dissociation (550 s) of pb10 chimeras.** Real-time SPR sensorgrams (resonance units, RU) obtained with 0 nM (control with the protein-free buffer) or 2.5 nM (saturating concentration) of pb10, pO and pNO. Analyzed data presented in **Figure 1d** include subtraction by the 0 nM curves. Note that pb10, pO and pNO associations induce different levels of response due to the proteins' different molecular weights. These differences were considered for the calculation of occupied binding sites in **Figure 1**, as described in the Methods.
